# *highroad* is induced by retinoids and clears mutant Rhodopsin-1 in *Drosophila* Retinitis Pigmentosa models

**DOI:** 10.1101/204131

**Authors:** Huai-Wei Huang, Brian Brown, Jaehoon Chung, Pedro M. Domingos, Hyung Don Ryoo

## Abstract

The light detecting protein, Rhodopsin, requires retinoid chromophores for their function. In vertebrates, retinoids also serve as signaling molecules, but whether these molecules similarly regulate gene expression in *Drosophila* remains unclear. Here, we report the identification of a retinoid-inducible gene in *Drosophila*, *highroad*, which is required for photoreceptors to clear folding-defective mutant Rhodopsin-1 proteins. Specifically, we identified *highroad* through an in vivo RNAi based genetic interaction screen with one such folding defective Rhodopsin-1 mutant, *ninaE*^*G69D*^. CRISPR-Cas9-mediated deletion of *highroad* results in the stabilization of folding-defective mutant Rhodopsin-1 proteins, and acceleration of the age-related retinal degeneration phenotype of *ninaE*^*G69D*^ mutants. Elevated *highroad* transcript levels are detected *ninaE*^*G69D*^ flies, and interestingly, deprivation of retinoids in the fly diet blocks this effect. Consistently, mutations in the retinoid transporter *santa maria* impairs the induction of *highroad* in *ninaE*^*G69D*^ flies. In cultured S2 cells, *highroad* expression is induced by retinoic acid treatment. These results indicate that cellular quality control mechanism against misfolded Rhodopsin-1 involves regulation of gene expression by retinoids.

## Introduction

As in many other metazoans, the *Drosophila* genome has several rhodopsin genes encoding the protein moiety Opsin, which becomes conjugated to a retinal chromophore. One such gene is *ninaE*, which encodes the seven transmembrane domain protein Rhodopsin-1 (Rh1) that binds the 11-cis-3-hydroxyretinal chromophore and is essential for outer photoreceptor function (Ahmad, 2006; O’Tousa, 1985; Zuker, 1985).

Certain types of mutations in Rhodopsins underlie Autosomal Dominant Retinitis Pigmentosa (ADRP), a disorder of age-related retinal degeneration (Dryja, 1990; Sung, 1991a). A majority of ADRP-causing alleles are missense mutations that impair the folding properties of the encoded protein, resulting in their instability (Sung, 1991b). This disease can be studied using a *Drosophila* model, where similar mutations in *ninaE* trigger age-related retinal degeneration (Colley, 1995; Galy, 2005; Kurada, 1995). For example, the *ninaE*^*G69D*^ allele results in the substitution of a transmembrane domain Gly to a negatively charged Asp residue in the first transmembrane domain of Rh1, and dominantly triggers age-related retinal degeneration in *Drosophila* (Colley, 1995). The product of the mutant *ninaE*^*G69D*^ (we henceforth refer to the encoded protein as Rh1^G69D^) fails to fold properly once synthesized in the endoplasmic reticulum (ER) resulting in activation of the Unfolded Protein Response (UPR) in the afflicted photoreceptors, thus affecting the course of retinal degeneration (Ryoo et al., 2007).

Recent studies point to the importance of ER and lysosomes as organelles involved in quality control against misfolded proteins. In the ER, a group of proteins are involved in the detection, retro-translocation and ubiquitination of misfolded peptides for proteasomal degradation in the cytoplasm, a process referred to as ER-Associated Degradation (ERAD) (Brodsky, 2012; Ruggiano, 2014). Proteomic studies have implicated more than 80 proteins forming an ERAD interaction network, which includes the ubiquitin ligase Hrd1 (Carvalho et al., 2006; Christianson, 2011; Denic et al., 2006). We had previously shown that overexpression of *hrd1* strongly delayed retinal degeneration in the *Drosophila ninaE*^*G69D*^ mutant, indicating that the photoreceptor’s ability to degrade misfolded rhodopsin is an important factor that affects the course of age-related retinal degeneration (Kang, 2009). In addition to ERAD, recent studies indicate that other types of misfolded proteins are trafficked out of the ER to be ultimately degraded in the lysosome, and rhodopsins are partly degraded through this mechanism (Chiang, 2012; Chincore, 2009; Satpute-Krishnan, 2014; Wang, 2014).

Rhodopsins fail to undergo proper maturation if photoreceptors are devoid of the chromophore retinal. Flies reared in a diet devoid of the chromophore, or its precursor vitamin A, have very low rhodopsin protein levels and show defects in photoreceptor electroretinogram profiles (Harris, 1977; Ozaki, 1993). Consistently, defects in rhodopsin maturation and photoreceptor defects can result from mutations that impair vitamin A transport and retinoid metabolism (Gu, 2004; Wang, 2007, 2005). In vertebrates, retinoids also have a second role as transcriptional regulators whose effects are mediated by the nuclear hormone receptor proteins (Mangelsdorf, 1995). Although previous studies reported that *Drosophila* that are deprived of vitamin A in the diet have altered levels of Opsin and fatty acid binding glycoproteins transcripts (Picking, 1996; Shim, 1997), the biological role and the mechanism of retinoid-mediated gene expression control in *Drosophila* remain unclear.

In this study, we report the identification of *highroad* (*hiro*), a gene that is required for mutant Rh1 degradation in *Drosophila* and also affects the course of age-related retinal degeneration. Furthermore, our data indicates that *hiro* transcript levels increase in *ninaE*^*G69D*^ mutant flies, and this is dependent on retinoid availability in vivo. These observations suggest that the degradation of mutant Rh1 is associated with retinoid-mediated gene expression control in *Drosophila*.

## Results

### Adult eye morphology based RNAi screen for genetic interactors of *ninaE*^*G69D*^

We had previously established a facile genetic assay system to assess cellular stress caused by *ninaE*^*69D*^ overexpression through the eye-specific *GMR* promoter (henceforth referred to as *GMR-Rh1*^*G69D*^). In these flies, Rh1^G69D^ is synthesized in early stages of eye development when the secretory pathway of the eye imaginal disc progenitor cells are not yet equipped to handle large loads of mutant Rh1^G69D^, resulting in adults emerging with malformed eyes (Kang, 2009; Kang, 2012; Also, see Figure S1). This phenotype can be attributed to increased misfolded proteins in the ER, since co-overexpression of the E3 ubiquitin ligase that promotes ERAD, *hrd1*, almost completely suppresses the external eye phenotype (Figure S1C-E).

To identify other factors involved in misfolded Rh1 quality control, we screened for RNAi lines that impaired the protective effects of *hrd1* overexpression against *GMR-Rh1*^*G69D*^ (Figure S1A). To make RNAi more efficient, we combined two eye-specific Gal4 drivers, *GMR-Gal4* and *ey-Gal4* to drive *UAS-RNAi* lines. In addition, we expressed *dicer-2* with the same Gal4 drivers in order to enhance the efficiency of RNAi knockdown (Dietzl, 2007). The screened RNAi lines included *Drosophila* homologs of mammalian genes with known roles in ERAD, or those that are found in proteins complexes with human HRD1, GP78 and their associated proteins (Christianson, 2011). We also included RNAi lines targeting annotated membrane proteases and carboxypeptidases in *Drosophila*. Many of these RNAi lines (e.g. *rhomboid-1, -5, -7, presenilin* and *spp*) caused eye phenotypes even in the absence of *GMR-Rh1*^*G69D*^ and *hrd1* overexpression, and those were excluded from further analysis.

As a positive control, we used an RNAi line that targets *hrd1*, which did not impair eye development when expressed alone, but aggravated the eyes of flies coexpressing *Rh1*^*G69D*^ and *hrd1* (Figure S1B). A number of other lines gave rise to phenotypes similar to *hrd1* knockdown. These included not only the lines that targeted *Drosophila* homologs of known ERAD genes, but also genes with no previous associations with ERAD, including CG32441, *asrij*, and CG3344 (Figure S1B).

### *Highroad* (*hiro*) is a gene required to reduce mutant Rh1 levels in photoreceptors

As a secondary assay for validation, we turned to the classical *ninaE*^*G69D*^ allele, with a mutation in the endogenous *ninaE* locus that results in very low levels of total Rh1 in newly eclosed adult flies (Colley, 1995; Kurada, 1995). Specifically, we expressed the RNAi lines of interest using *Rh1-Gal4* in photoreceptors of *ninaE*^*G69D*^/+ flies, and examined total Rh1 protein levels in the fly head extracts by western blot. Only one RNAi line (VDRC 110402) had a dramatic effect on the Rh1 levels in *ninaE* ^*G69D*^/+ flies, almost fully restoring Rh1 levels in *ninaE*^*G69D*^/+ background to wild type levels (Figure 1A, B). This line targets a previously uncharacterized carboxypeptidase, *CG3344*, homologous to a mammalian protein known as ‘retinoid-inducible serine carboxy peptidase’ or ‘serine carboxy peptidase 1’ (SCPEP1) (Chen, 2001). The RNAi line targeting *CG3344* was included in our screen due to its remote homology to CPVL carboxypeptidase, a reported member of the Hrd1 interaction network (Christianson, 2011), but it is not an ortholog of that protein. Based on the loss of function phenotype, we henceforth refer to this gene as HIGH RhOdopsin Accelerated Degeneration or *highroad* (*hiro*).

**Figure 1:**
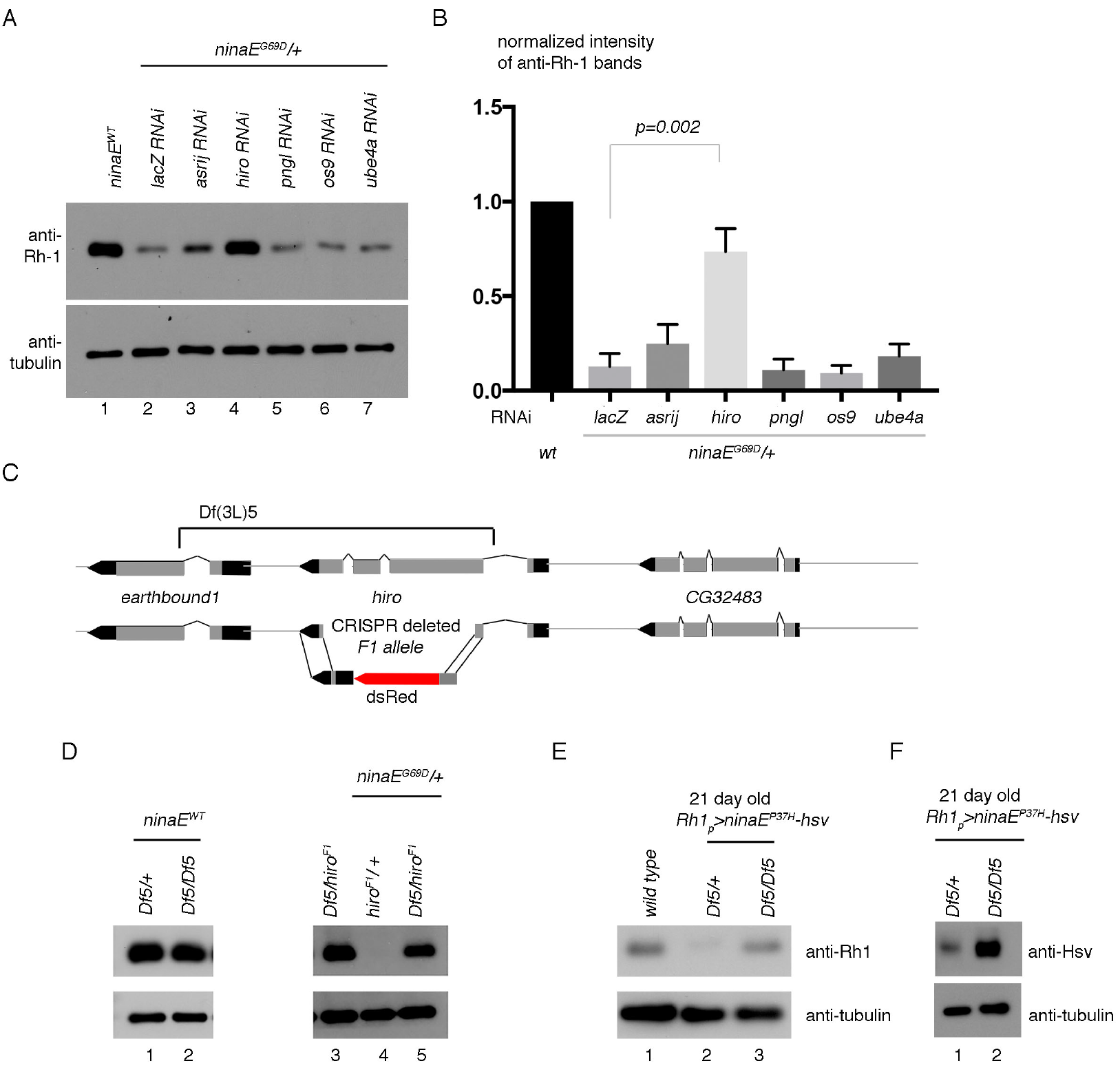
*hiro* is required for photoreceptors to reduce mutant Rh-1 levels. (A) Shown are western blots of adult head extracts with the indicated antibodies. Compared to *ninaE* wild type heads (lane 1), *ninaE*^*G69D*^/+ mutant head extracts contain dramatically reduced Rh1 protein levels (lane 2). Lanes 2-7 are from *ninaE*^*G69D*^/+ samples in which the indicated genes were knocked down with *Rh1-Gal4* driver specifically in photoreceptors. (B) Quantification of the band intensities shown in (A). (C) A schematic diagram of the *hiro* locus. The bar above indicates sequences deleted in *Df(3L)5*. Below is a schematic diagram of the *hiro*^*F1*^ allele that was generated through CRISPR-Cas9. (D, E) Validation of *hiro RNAi* phenotype through classical alleles. Anti-Rh1 (top) and anti-beta tubulin (bottom) western blot from adult fly head extracts of the indicated genotypes. The left panel shows results in the wild type background and the right panel in the *ninaE*^*G69D*^/+ background. (E, F) Anti-Rh1 western blots with *ninaE*^*G69D*^ were performed with 0-5 day old adult fly head extracts (E), and with *Rh1*_*p*_>*ninaE*^*P37H*^, fly head extracts with 21 day old fly samples (F).

To further validate the RNAi result from the screen, we employed two *hiro* mutant alleles. 1) The *Df(3L)5* deficiency, in which *hiro* and its neighboring gene *earthbound1* are deleted (Benchabane, 2011) and 2) CRISPR-Cas9 mediated deletion of the *hiro* locus (Figure 1C; also Experimental Procedures), which we refer to henceforth as *hiro*^*F1*^. *highroad*^*F1*^ −/− or *hiro*^*F1*^/*Df(3L)5* flies were viable and did not exhibit any obvious morphological phenotype. In *ninaE wild type* background (*w*^*1118*^), the loss of *highroad* in the *Df(3L)5* −/− or *Df(3L)5*/*hiro*^*F1*^ backgrounds did not change Rh1 protein levels (Figure 1D, lanes 1, 2 and 3). However, in the *ninaE*^*G69D*^/+ background, loss of *hiro* in *Df(3L)5*/*highroad*^*F1*^ resulted in the recovery of Rh1 levels comparable to that of wild type flies (Figure 1D, lane 5). Together, these results validate that *hiro* is genetically required to reduce Rh1 levels in *ninaE*^*G69D*^/+ mutants.

### *highroad* affects the levels of other Rh1 mutant alleles

In humans, the most widespread *rhodopsin* allele associated with ADRP is the *P23H* mutation, which results in improperly folded Rhodopsin that undergoes degradation in the ER and the lysosome (Chiang, 2012; Dryja, 1990; Liu, 1996). This mutation has been modeled in *Drosophila* by substituting Proline 37 in Rh1 to a Histidine residue, and expressing this allele tagged with an HSV-epitope in photoreceptors with the *ninaE* promoter in transgenic flies (Galy, 2005). Unlike the *G69D* mutants, we found that Rh1 levels were not significantly decreased in newly eclosed *P37H* flies (data not shown). However, *P37H* flies showed an age-dependent decrease in Rh1 levels, with overall Rh1 level significantly lower than wild type controls at 21 days post-eclosion (Figure 1E, lane 2). Similar to results with the *G69D* mutants, Rh1 levels in *P37H* mutants were almost restored to wild type levels in the *Df(3L)5* −/− background (Figure 1E, lane 3). Consistently, western blotting with anti-HSV to specifically detect *P37H* mutant protein (rather than total Rh1) showed higher P37H-HSV in the *Df(3L)5* −/− background (Figure 1F). These results show that the effect of *highroad* on Rh1 is not specific to the *ninaE*^*G69D*^ allele, but is applicable to other disease relevant *ninaE* mutant alleles.

### Epitope tagged Hiro co-localize with a lysosomal marker

To determine the sub-cellular localization of Hiro protein, we generated a V5-epitope tagged *uas-hiro* line and expressed it in various tissues. As a lysosome marker, we used anti-ATP6V1B1 antibody that labels subcellular structures in cells with high lysosome content, such as the 3^rd^ instar larval fat body cells (Figure 2). The V5 epitope tag was detected in these cells in a pattern indistinguishable from that of ATP6V1Ba. On the other hand, ER as visualized with anti-Hsc3 antibody gave a non-overlapping pattern with that of Hiro-V5 (Figure 2). The observation is consistent with the reported lysosomal localization of the mammalian homolog SCPEP1 (Kollmann, 2009; Lee, 2006).

**Figure 2:**
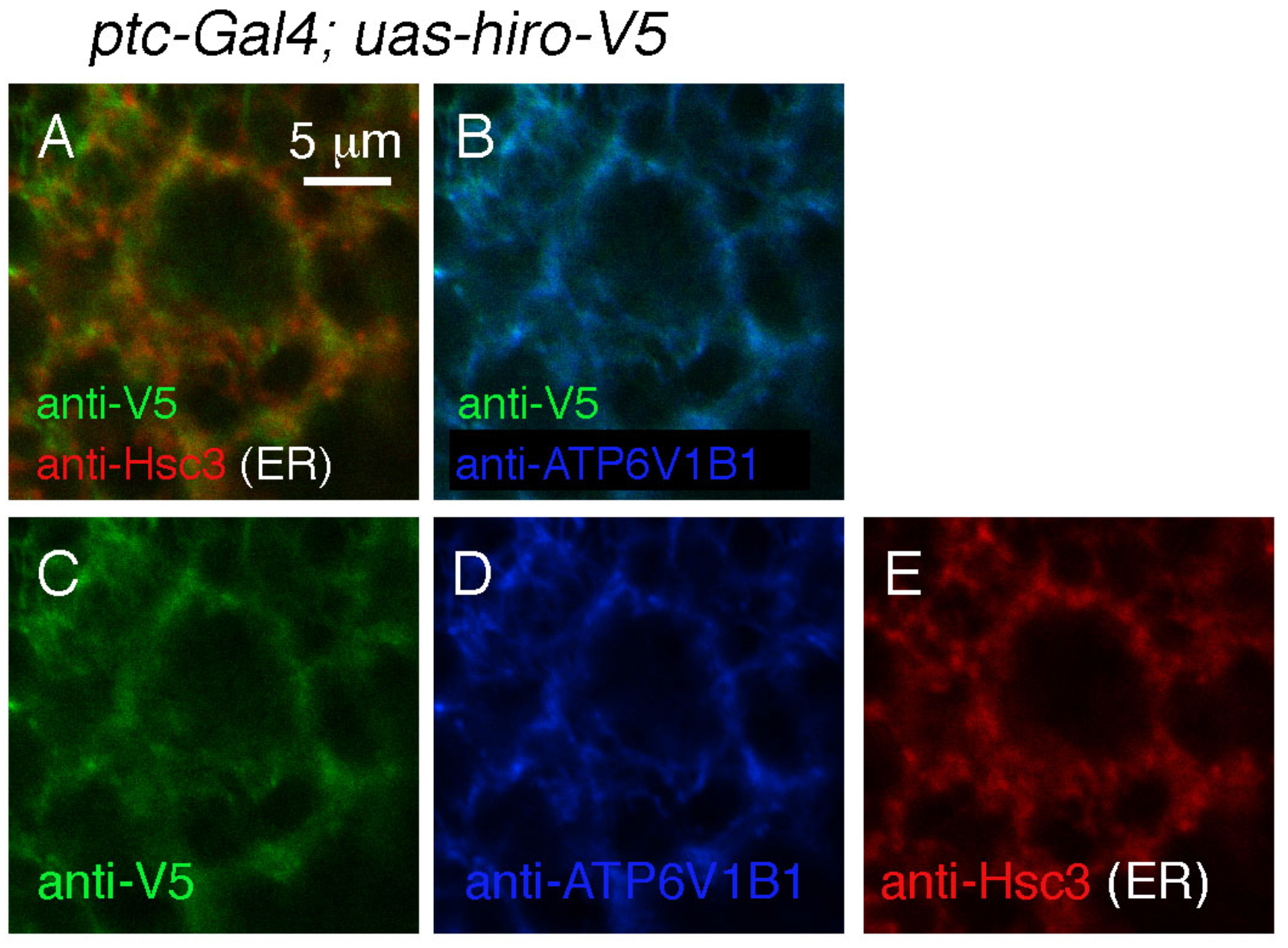
Subcellular localization of Hiro. V5 epitope tagged *hiro* was expressed with the *ptc-Gal4* driver. Shown are fat body cells labeled with anti-V5 (green), anti-Hsc3 that marks the ER (red) and anti-ATP6V1B1 (blue) that marks the lysosome. (A) An image with anti-V5 and anti-Hsc3 channels merged shows distinctively red and green signals, indicating that the Hsc3 localization pattern is different from that of anti-V5. (B) A merged image with anti-V5 and anti-ATP6V1B1. The signal appears mostly as light blue indicative of co-localization. Individual channels of images are shown in panels, C, D and E. The scale bar in A applies to all panels.

### *ninaE*^*G69D*^/+ show accelerated retinal degeneration in the absence of *hiro*

Since our data indicated that *hiro* mutants fail to properly regulate Rh1 levels, we decided to examine if this affected the age-related retinal degeneration of *ninaE*^*G69D*^/+ flies. We tested this using the *Rh1-GFP* reporter that allows the examination of photoreceptor integrity in live flies over a time course of 30 days. A majority of *ninaE*^*G69D*^/+ flies showed an organized trapezoidal pattern of Rh1-GFP up to 20 days, before showing signs of degeneration (Figure 3A). In the *hiro*^*F1*^-/- background, *ninaE*^*G69D*^/+ flies showed earlier signs of retinal degeneration (p<0.0001, Chi square 178.1, when compared to *ninaE*^*G69D*^/+), with more than 75% of flies showing disorganized Rh1-GFP patterns at 20 days old (Figure 3A). The control *hiro* -/- flies in the *ninaE wild type* background showed no signs of retinal degeneration at these time points. We independently validated these results by dissecting representative fly eyes (Figure 3B). In wild type flies, Phalloiding labeling marks the rhabdomeres in each ommatidia, revealing the typical trapezoidal pattern of the photoreceptors (Figure 3A). In *ninaE*^*G69D*^/+ flies up to 20 days old, this pattern is maintained in most dissected flies (Figures 3B). On the other hand, *ninaE*^*G69D*^/+; *hiro*^*F1*^ −/− fly eyes showed the absence of the trapezoidal rhabdomere arrangement by day 20, indicative of retinal degeneration (Figure 3B, lower right most panel). Thus, these results indicate that *hiro* has a protective effect against retinal degeneration in *ninaE*^*G69D*^/+ flies.

**Figure 3:**
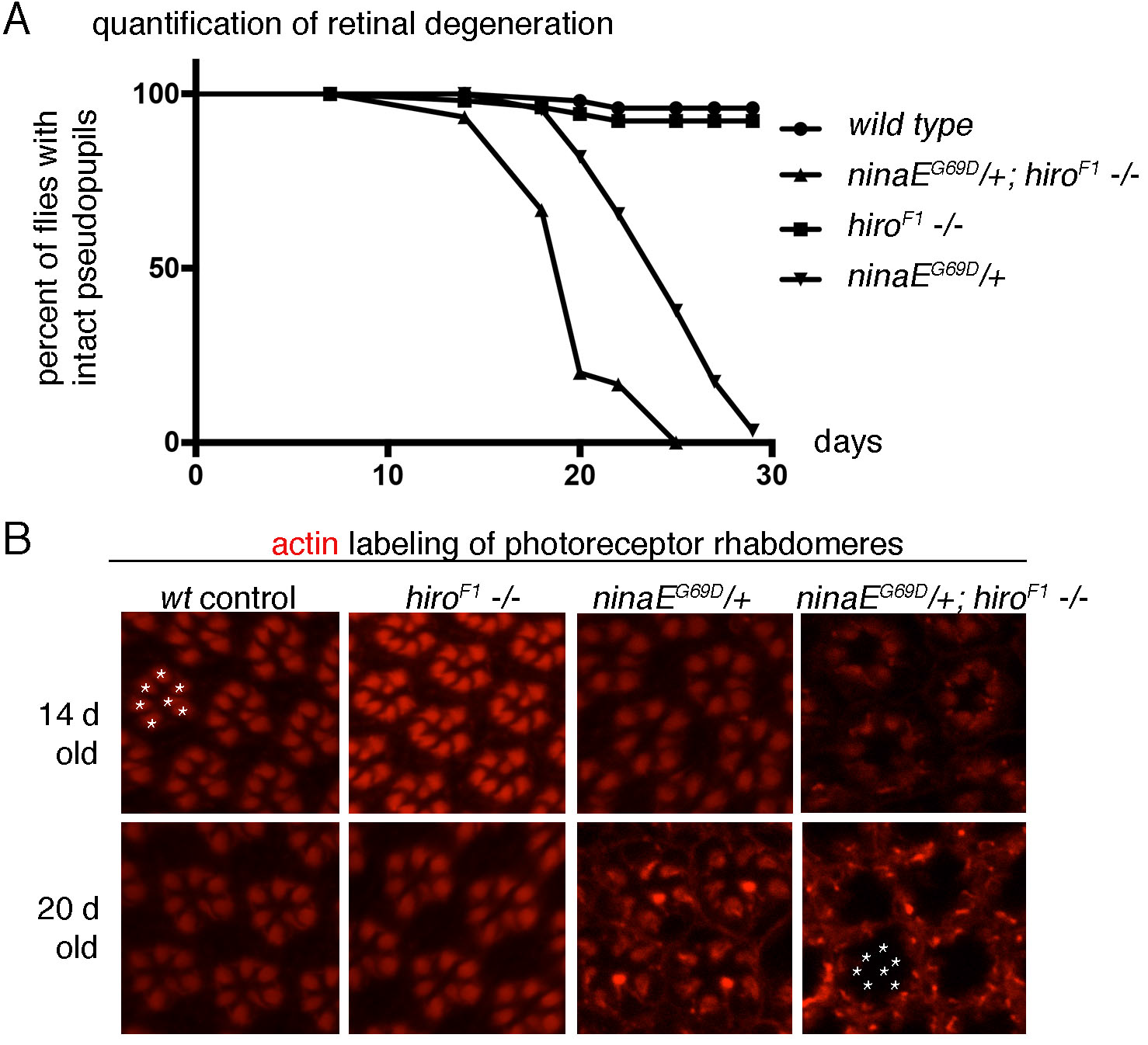
Loss of *highroad* accelerates the course of age-related retinal degeneration in *ninaE*^*G69D*^/+ flies. (A) Retinal degeneration was assessed in live flies containing Rh1-GFP to visualize photoreceptors, quantified and exhibited as a graph. At day 20, most *ninaE*^*G69D*^/+ control flies still maintained a regular pattern of Rh1-GFP, whereas most of the *ninaE*^*G69D*^/+; *hiro* −/− eyes showed disorganized patterns indicate of retinal degeneration. (B) Representative images of dissected ommatidia labeled with Phalloidin that labels rhabdomeres of photoreceptors (red). The genotypes are indicated on top of each panels. Intact ommatidia show trapezoidal arrangement of 7 photoreceptor rhabdomeres (one example marked with asterisks on the upper left panel). On the other hand, 20 day old *ninaE*^*G69D*^, *hiro* −/− eyes (lower right most panel) lack the trapezoidal pattern of Phalloiding labeling. The asterisks in that panel indicate the expected positions of Phalloiding-positive rhabdomeres in an ommatidium, which is lacking in this genotype.

### *hiro* is inducible by retinoids

The mammalian homolog of *hiro* is a gene whose expression is induced by retinoic acids (Chen, 2001). To test whether *Drosophila hiro* similarly responds to retinoic acids, we challenged *Drosophila* S2 cells with commercially available all-trans retinoic acid. We found that 20 minutes of retinoic acid treatment resulted in increased *hiro* transcript levels as evidenced by semi-quantitative RT-PCR as well as qPCR analyses (Figure 4A, B).

**Figure 4:**
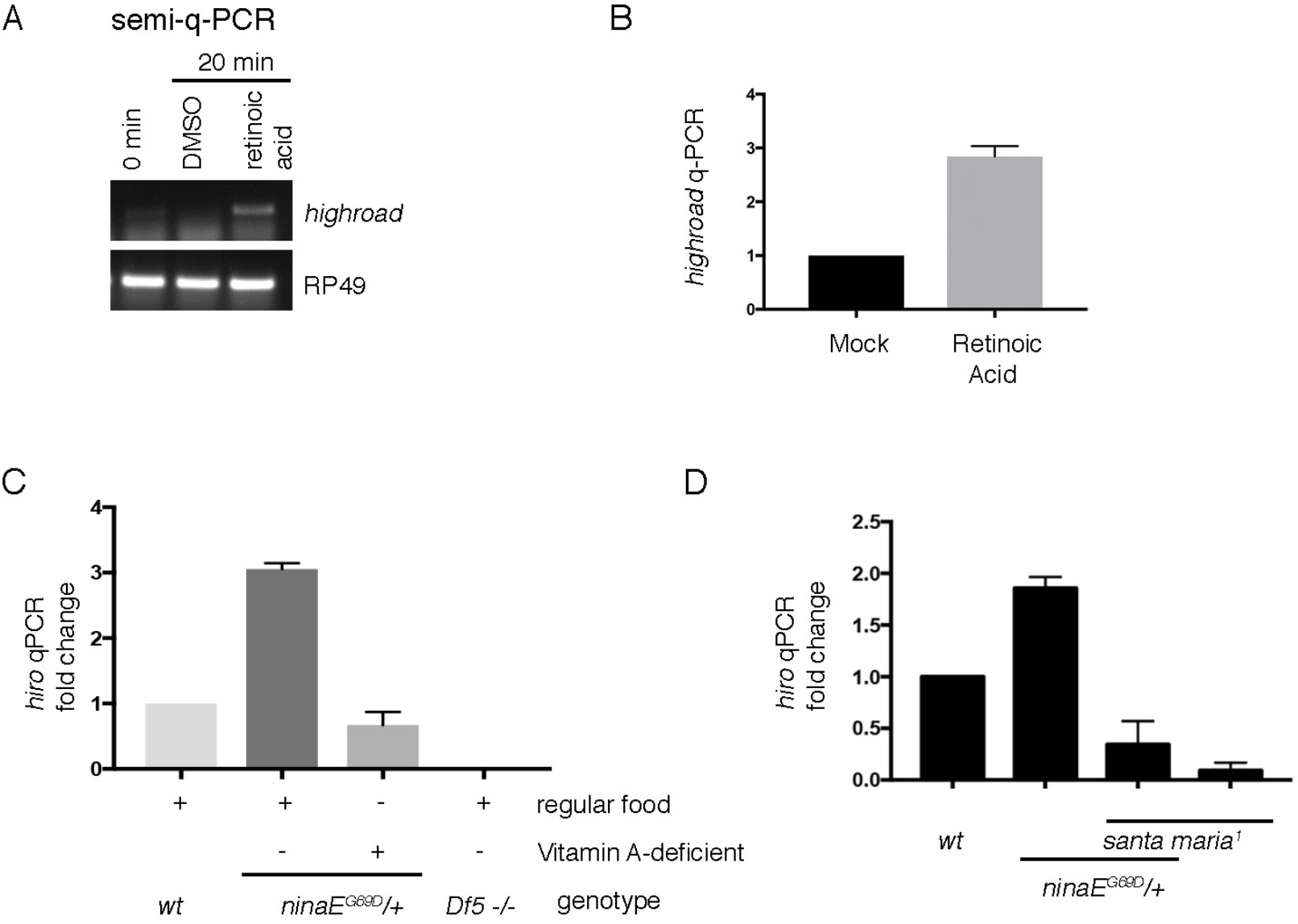
*hiro* transcript levels are regulated by retinoids. (A) Semi-quantitative RT-PCR of *hiro* (top) and the control RP49 (bottom) from cultured *Drosophila* S2 cells with or without retinoic acid treatment. (B) qPCR of *hiro* from S2 cells treated with DMSO (left) as control or with retinoic acids (right). (C) qPCR of *hiro* from fly head extracts of indicated genotypes. Flies were either raised in regular cornmeal medium or in vitamin A deficient food. (D) qPCR analysis of *hiro* induction in wild type (wt) and in *santa maria*^*1*^ mutants that impair retinoid transport to the nervous system (right).

qPCR analysis of *hiro* from adult *Drosophila* heads also detected higher signals from *ninaE*^*G69D*^/+ samples, as compared to the wild type controls (Figure 4C, lanes 1 and 2). To determine if such *hiro* induction in fly heads is due to retinoid-induced gene expression, we repeated the quantitative RT-PCR analysis under retinoid deprivation conditions. Since metazoans require dietary vitamin A to produce retinoids and related metabolites, one way to achieve retinoid deprivation is to rear flies in vitamin A deficient food (Blomhoff, 1990; Harris, 1977; Ozaki, 1993). We found that such conditions blocked the increase in *hiro* transcripts in *ninaE*^*G69D*^/+ heads (Figure 4C lane 3). To independently validate the role of retinoids in the regulation of *hiro* transcript levels, we employed genetic methods for retinoid deprivation in the eye. We specifically utilized the *santa maria* mutant that has impaired vitamin A/β-carotene transport to the photoreceptors (Wang, 2007). Similar to the results obtained with flies reared under vitamin A deficient food, loss of *santa maria* impaired *hiro* induction in the *ninaE*^*G69D*^/+ fly heads (Figure 4D). These results support an unexpected idea that retinoids regulate gene expression in *Drosophila*, and one such regulated gene *hiro* is involved in the clearance of mutant Rh1.

## Discussion

As rhodopsin misfolding underlies many cases of Autosomal Dominant Retinitis Pigmentosa (ADRP), boosting a cell's ability to clear misfolded rhodopsins may prove to be an effective way to develop therapies against ADRP. Related to this idea, much focus had been placed on the mediators of ER-Associated Degradation (ERAD), including our own previous work showing that overexpression *hrd1*, a central ubiquitin ligase for ERAD, can delay the course of age-related retinal degeneration in a *Drosophila* model for ADRP (Kang, 2009). We began this study with an RNAi screen to identify factors involved in the degradation of mutant Rh1. Surprisingly, the strongest effect was observed after knocking down an uncharacterized gene that we now name *hiro*, instead of the known ERAD components. The lack of strong effects with ERAD component knockdown is consistent with what we had reported previously (Kang, 2009). The loss of *hiro* not only leads to increased levels of Rh1, but also accelerates the course of retinal degeneration in a *Drosophila* model for ADRP.

We found that *hiro* transcripts are induced by retinoic acids in cultured S2 cells, and the increase in *hiro* in response to *ninaE*^*G69D*^/+ requires dietary vitamin A and retinoid transporter gene activity in vivo. It is noteworthy that while *Drosophila* is a popular model organism for studying various cell signaling pathways, retinoic acid signaling had not been examined carefully in this organism. This is partly because unlike vertebrates, manipulating retinoid levels does not result in gross morphological changes during development. In fact, the effect of retinoic acid deprivation in *Drosophila* has been largely limited to two phenotypes: One is a role of retinoids and its metabolites in halting developmental progression in response to X-ray irradiation during *Drosophila* larval development (Halme, 2010). Whether that effect is due to retinoic acid-mediated gene expression changes, or alternatively, through metabolic effects of retinoids remains unknown. The other phenotype associated with retinoid deprivation is the inability of photoreceptors to respond to light (Gu, 2004; Harris, 1977; Kiefer, 2002; Ozaki, 1993; Wang, 2007, 2005). This phenotype has been largely attributed to retinoid’s role as a chromophore for rhodopsins. However, at least two studies have reported changes in gene expression under conditions of retinoid deprivation *Drosophila* (Picking, 1996; Shim, 1997). The evidence we present in this study further supports the idea that retinoic acids can change gene expression in *Drosophila*, and that one such responsive gene is involved in mutant Rh1 quality control.

As normally folded Rh1 is bound to an 11-cis hydroxylretinal (Ahmad, 2006), our results raise the possibility that these retinoids are released to the cytoplasm of photoreceptors with Rh1 folding mutations, thereby serving as a second messenger to instruct the expression of quality control genes that clear misfolded Rh1. Such a speculative idea awaits more in depth examination in future studies, along with a search for the transcription factors that directly bind retinoids for gene expression regulation in *Drosophila*.

## Acknowledgements

We thank J. Wildonger and M.J. Kang for technical advice, and A. Galy, C. Montell, C. Desplan, VDRC and Bloomington stock centers for fly strains and antibody. This work was supported by the NIH grant, R01 EY020866 to H.D.R. PMD was supported by grants LISB0A-01-0145-FEDER-007660, FCT-ANR/NEU-NMC/0006/2013, PTDC/NEU-NMC/2459/2014 and IF/00697/2014.

## Experimental Procedures

### Fly Genetics

The following flies used in this study had been reported previously: *GMR-Rh1*^*G69D*^ (Kang, 2009) *ninaE*^*G69D*^ (Colley, 1995), *Rh1-GFP* (Pichaud and Desplan, 2001), *Rh1-Gal4* (Mollereau et al., 2000), *santa maria*^*1*^ (Wang, 2007), *Df(3L)5*(Benchabane, 2011), *Rh1*_*p*_>*ninaE*^*P37H*^ (Galy, 2005), *uas-dicer2* (Dietzl, 2007).

RNAi screen: Inverted repeat UAS (*UAS-RNAi*) lines from the Vienna Stock Center were crossed to the female virgins of the following genotype: *GMR-Gal4; ey-Gal4, GMR-Rh1*^*G69D*^, *uas-hrd1*/*CyO*; *uas-dicer2*. As a control, we also crossed the *UAS-RNAi* lines to flies that lack the *GMR-Rh-1*^*G69D*^ and *uas-hrd1*. The genotypes of those control flies were: *GMR-Gal4; ey-Gal4/CyO; uas-dicer2.* To validate the hits with the *ninaE*^*G69D*^ endogenous allele, *UAS- RNAi* lines were crossed to the virgin females of the genotype; *Rh1-Gal4; uas-dicer2; ninaE*^*G69D*^/*TM6B*. Non-TM6B progeny were collected to generate head extracts for anti-Rh1 western blot analysis.

Generation of *highroad*^*F1*^: To generate *highroad* deletion mutants, we followed the homology directed repair CRISPR-Cas9 protocol available from flycrispr.molbio.wisc.edu. The following sequences were subcloned into pU6-Bbsl-chiRNA to generate guide RNA plasmids.

5’-AATATACTCTCAGCACGCAA-3’
5’-GCGACATGGCCGCGGGATTAT-3’

Donor plasmids for homology-directed repair were generated by subcloning 1.0 and 1.3 kb homology arms distal to the 5’ and 3’ CRISPR-Cas9 cut sites respectively were subcloned into the pHD-DsRed-attP vector. The donor plasmid and the guide RNA plasmids were injected into *nanos-Cas9* embryos (by Best Gene Inc.). The resulting mosaic adult survivors were crossed to *w*^*1118*^ flies and screened for dsRed positive progeny. The deletion of CG3344 was further confirmed through genomic PCR.

### Immunofluorescence and Western Blots

Standard protocols were followed for western blots and whole mount immuno-labeling. The following primary antibodies were used in this study: Mouse monoclonal 4C5 anti-Rh1 (Developmental Studies Hybridoma Bank, used at 1:500 for whole mount, and 1:5000 for western blots), rabbit anti-Rh1 polyclonal antibody (Mitra, 2011), anti-β tubulin antibody (Covance #MMS-410P), rabbit anti-ATP6V1Ba antibody (abgent #AP11538C, used at 1:10), guinea pig anti-Hsc3 (Ryoo et al., 2007) (used at 1:50). Rhodamine-conjugated Phalloidin (Molecular Probes cat #R415) was used to detect rhabdomeres in whole mount stainings.

### RT-PCR

qRT-PCR was performed using Power SYBR green master mix kit (Thermo Fisher) with the following primer pairs:

qCG3344 F GGAATTGGAACTGAAACACTGG
qCG3344 R TCCTGACATCCACAAATCCC

For semi-quantitative RT-PCR, the following oligos were used to amplify a 280 bp product:

Semi-qCG3344 F2 TCGGTGTCTCTTGGAATTGG
Semi-qCG3344 R2 AAGTTTCCGTATCCCGTTGAG

### Retinal degeneration assay

All retinal degeneration assays were done in the *cn*, *bw* −/− background to eliminate eye pigments, which could otherwise affect the course of retinal degeneration. To generate the retinal degeneration graph in Figure 3A, green fluorescent pseudopupils appearing from *Rh1-GFP* was analyzed in live flies. A clear trapezoidal pattern of GFP under low magnification microscope was interpreted as having intact photoreceptors, while the disappearance of the trapezoidal pattern was construed as a sign of retinal degeneration. The number of flies analyzed of each genotype is as follows: *w*^*1118*^ (*n*=*52*), *hiro*^*F1*^ (*n*=*54*), *ninaE*^*G69D*^ (*n*=*49*), *ninaE*^*G69D*^; *hiro*^*F1*^ −/− (*n*=*31*). The results from the pseudopupil assay were further substantiated through Phalloidin labeling of dissected fly eyes as shown in Figure 3B.

### Statistics

We quantified anti-Rh1 band intensities in Figure 1 by measuring average pixel intensities of western blot bands using Image J and normalizing them to anti-β tubulin bands. The data are representative of the average of three independent experiments and *p* values were calculated through a paired paired t-test. For retinal degeneration analysis, we applied the Log-rank (Mantel-Cox) test.

## Supplemental Information

**Figure S1:**
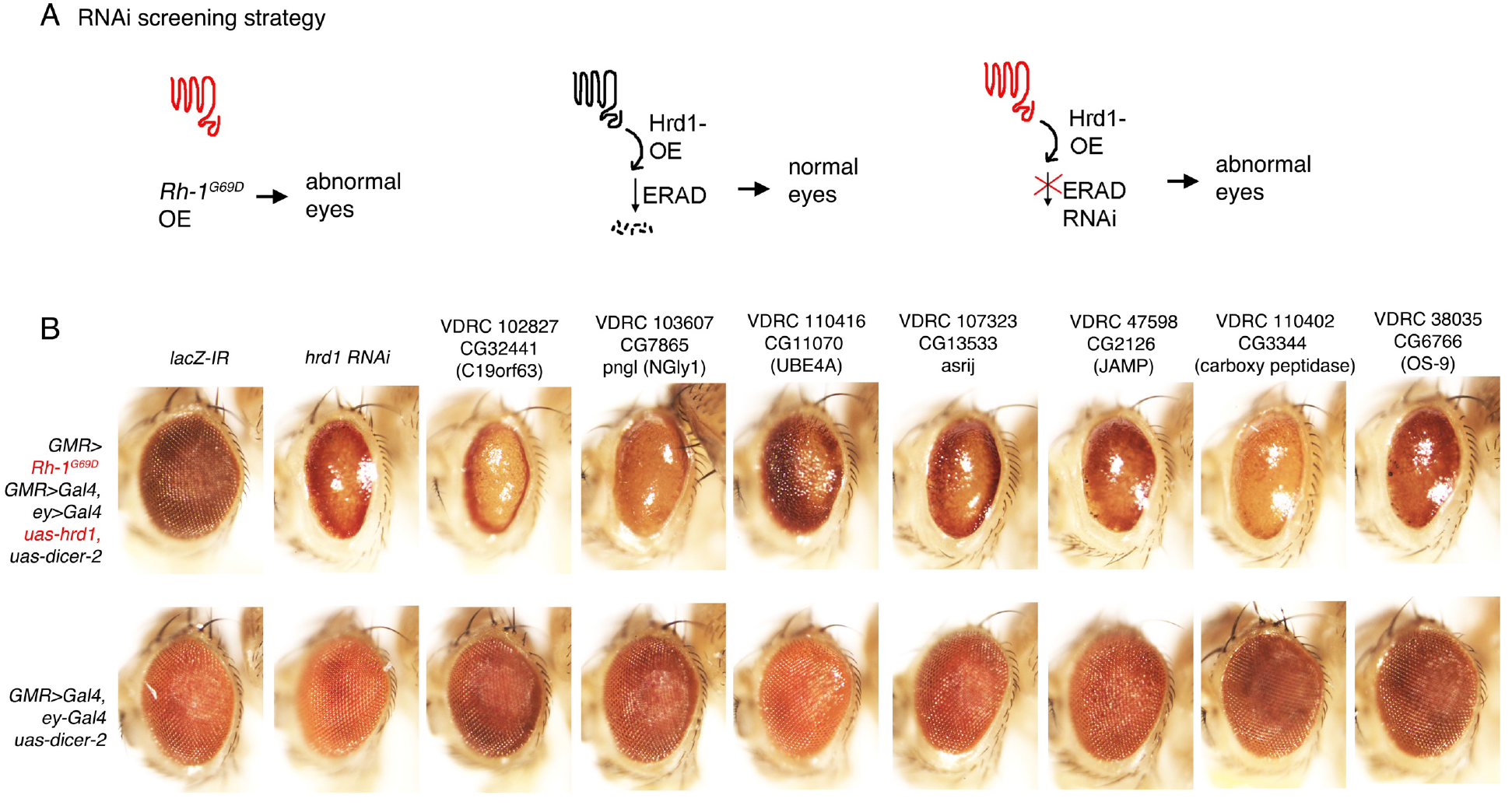
*Drosophila* eye based RNAi screen for factors involved in ER-Associated Degradation (ERAD) of Rh-1^G69D^. (A) Schematic diagram of the screening strategy. Overexpression of mutant Rh-1^G69D^ during eye development results in abnormal adult eyes due to proteotoxicity, which is suppressed by concurrent overexpression of *hrd1*, a ubiquitin ligase involved in ERAD. RNAi-mediated knockdown of genes essential of ERAD in this background is expected to impair the protective function of *hrd1*, thereby causing eye abnormalities. (B) The upper panels show fly eyes that overexpress *Rh-1*^*G69D*^ and *hrd1*, along with RNAi lines that target the indicated genes. The lower panels are genetic controls to show that knock down of the indicated genes do not cause developmental defects on their own. The relevant genotypes used are indicated in the left margin. *lacZ* RNAi was used as negative control and *hrd1* knockdown was used as a positive control. Only hits that generated phenotypes similar to *hrd1* knockdown are shown. The mammalian homologs of the knocked down genes are indicated in parentheses.

